# From code to natural language: MErlin – a multi-omics toolkit for bacterial epigenomics delivered as Claude agent skill

**DOI:** 10.64898/2026.09.02.748773

**Authors:** Iacopo Passeri, Solène Pety, Michele Giovannini, Marco Fondi, Alessio Mengoni, Elena Perrin

## Abstract

Interpreting a bacterial methylome is a multi-omics problem. It requires integrating modified-base calls with genome annotation, motif inventories, methyltransferase genotypes, transcript abundance, replichore position and, increasingly, chromosome conformation. These data types are commonly generated in incompatible formats, use inconsistent sequence and gene identifiers, and originate from different analytical workflows. Relevant algorithms are available but assembling them into a coherent and statistically defensible analysis remains a substantial data-integration and interface problem.

We present MErlin (Methylation-driven Expression & Regulation Linkage in Interacting Nuclear-domains), a multi-omics toolkit comprising seventeen composable modules, from basecalled modBAM files to ranked gene-level evidence and a self-contained HTML report. MErlin is distributed both as a conventional Python package and as an agent skill: a structured, version-controlled layer of procedural knowledge that enables a compatible large language model (LLM) assistant to select and operate the audited package without generating a new analysis implementation for each request. This design treats natural language as an interface to fixed analytical operations rather than as a substitute for tested scientific software. The skill encodes module-selection rules, mandatory preflight checks, questions that require human input, design-to-inference constraints, and interpretation guidance. We describe MErlin’s architecture and statistics, validate it against a synthetic dataset with planted ground truth, and illustrate the conversational interface on a real methylome–transcriptome comparison in *Pseudoalteromonas haloplanktis* TAC125.

MErlin is open source and available at https://github.com/IacopoPasseri/MErlin.

## 1. Introduction

For most of the twentieth century, DNA methylation in bacteria was primarily understood as the self/non-self marking arm of restriction–modification (R-M) systems [1]. That view is now recognised as incomplete in two respects. First, R-M systems are more than nucleases: their genomic distribution tracks mobile-element content, and by determining which incoming DNA survives, they moderate horizontal gene flow rather than simply blocking it [2]. Second, and more consequential for what follows, methylation is a signal read by proteins other than restriction enzymes [3]. Prokaryotic adenine and cytosine methylation contribute to mismatch repair, to the timing of replication initiation, to the cell-cycle-coupled program of *Caulobacter* and other *alphaproteobacteria* [3,4], to the heritable switching of phase-variable regulons through phasevarions [5], and to transcriptional modulation at promoters where a methylase and a regulator compete for the same DNA site [6,7]. Single-molecule sequencing moved this from a set of case studies towards a survey discipline: SMRT kinetics [8] and, later, nanopore signal-level modification calling [9,10] base-resolution mapping of 6mA, 4mC and 5mC across a whole bacterial genome routine, a revealing that most sequenced prokaryotes carry at least one methylated motif, frequently with no cognate restriction enzyme [11,12].

The phenotypic weight of DNA methylation is not predictable from the motif itself; establishing it generally requires several data layers at once. For instance, in *Clostridioides difficile*, a species-conserved methyltransferase was shown to be required for normal sporulation and for colonization in vivo, an attribution that rested jointly on comparative methylomes, transcriptome, targeted knockout and an animal model together, and that no single approach would have supported [13]. Such conserved methyltransferases are widespread enough across taxa to make this a general question rather than a curiosity of one pathogen [14]. Evolutionary designs offer a complementary route: in the plant symbiont *Sinorhizobium meliloti*, we recently crossed methylome with genomic organization across multiple strains to define a pan-epigenome and genomic context preferences for selected DNA methylation motifs [15]. Holding the genome nearly constant while the methyltransferase complement varies is one of the two contrasts, we argue to be most informative, and the context-preference result illustrates the point made throughout this paper: the finding emerges not from any single analysis, but from motif, positional and conformational layers examined together. The consequence is that the interesting questions in bacterial epigenomics are almost all relational.

A methylome on its own is a list of coordinates. It acquires meaning only when set against something else:

- the annotation, to ask which features are methylated and where within them — a mark in a promoter or operator, where it can occlude or be occluded by a regulator, invites a different mechanistic hypothesis from marks distributed evenly through a coding sequence;
- the motif inventory and the methyltransferase complement, to ask which system is responsible and whether it is orphan or R–M-coupled [11]; the distinction biologically meaningful, because a solitary methyltransferase has no defensive partner to account for its retention and is thus a more plausible candidate for a regulatory role [14];
- a second condition or a knockout, to ask what changes — and, when the knockout is of the methyltransferase itself, to move from a correlation to a manipulable cause;
- the transcriptome, to ask whether methylation and expression co-vary, and with what sign;
- the replichore geometry, because methylation state near the origin is confounded with replication timing in any growing culture: sites behind the fork are transiently hemimethylated until the maintenance methyltransferase catches up, so an origin-proximal deficit can be a property of the culture rather than of regulation [4];
- the reads themselves, because a site at 50% methylation in a population average may reflect every cell carrying one methylated strand or half the cells fully methylated and half not — the difference between a replication intermediate and an epigenetic switch, and one that is visible only at single-molecule resolution [5,16];
- the mobile fraction of the genome, because prophages, plasmids, integrative elements and genomic islands frequently carry their own methyltransferases and are the substrate on which R–M systems act, so their methylation state need not resemble that of the core genome [2];
- the chromosome conformation, because chromosomal interaction domains organize bacterial genomes at the scale at which co-methylation might plausibly be coordinated [17,18].

In our view, none of these comparisons is optional. A claim that a given mark is regulatory requires four things at once: (i) that the mark lies in a position where a DNA-binding protein could plausibly be affected by it, typically a promoter or operator; (ii) that its methylation state changes with growth condition or upon deletion of the responsible methyltransferase; (iii) that transcription of the associated gene changes in the same comparison; and (iv) that trivial explanations — replication timing, local sequence composition, uneven coverage — have been ruled out. Each of these lives in a different data layer, which is why bacterial epigenomics is an integration problem before it is a statistical one. Hemimethylation illustrates why the layers should be considered jointly rather than in sequence. It simultaneously the principal confounder of any methylation measurement taken in a growing culture and, in cell-cycle-coupled systems such as CcrM/GANTC in alphaproteobacteria, one of the most informative signals available [3,4]. A single read-level measurement both, and they are separable only against replichore position: hemimethylation that decays with distance behind the replication fork is a property of replication, whereas hemimethylation that persists independently of position is a candidate for regulation. Neither phase nor space can make that distinction alone; their conjunction can.

Each of these joins is individually tractable but together they are a data-engineering project. Files arrive as bedMethyl, modification GFF3 or modBAM with MM/ML tags; as Prokka [19] or NCBI GFF3; as a DESeq2 [20] results table with an undocumented sign convention; as a .cool or .mcool matrix at a resolution that does not match the gene scale. They use different sequence-identifier namespaces and were produced by different people, sometimes years apart. The analyst who needs to join them is, in the median case, a microbiologist who designed the experiment and now needs an answer and not a software engineer.

This difficulty is not peculiar to bacterial epigenomics. In the largest survey of its kind, 89% of 704 National Science Foundation (NSF) Biology principal investigators reported an unmet need for training in the integration of multiple data types — the highest-ranked deficiency, above data management and above scaling to HPC, while hardware and storage ranked lowest [21]. The practical bottleneck is therefore not computation alone, but the work required to make heterogeneous data products interoperable and to determine what an integrated result is allowed to mean.

In multi-omics workflows, the heterogeneity of data products becomes consequential for one specific reason: an integration step can fail without producing an explicit software error. Every integration of the kind described above reduces, in the end, to one of two joins — on a sequence identifier (seqid) or on a gene identifier (locus_tag) — and both fail silently. If the FASTA reads NZ_CP012345.1 and the Prokka GFF3 readsNZ_CP012345, an inner join returns zero rows: no exception, no warning from pandas, from bedtools, or from any of the obvious ways would write the operation by hand. The output is a table of zeros, which is precisely - and indistinguishably-what a genome with no methylation would produce. This is a particularly insidious class of scientific software error: it produces no stack trace, yield apparently valid output, and may therefore acquire a biological interpretation. It is also preventable, provided identifier compatibility is verified before analysis rather than assumed, and provided that verification is an explicit operation rather than an implicit expectation of user expertise.

The longstanding response to usability barriers in computational biology has been to place graphical or web-based interfaces over command-line tools. Galaxy is a prominent and successful example [22]. The web-form model, however, imposes its own costs: a fixed set of exposed parameters, a deployment to maintain, and a mapping between the user’s actual question and the available boxes that the user must perform themselves. The user still has to know which tool they want before they can find its form.

A different possibility has opened over the last two years, largely outside of bioinformatics. Large language model assistants have become an ordinary part of professional practice, used routinely, by people with no particular interest in the technology, for tasks previously done by hand. In parallel, these assistants acquired the ability to act: to read files the user provides, to run code, to inspect the result and iterate. The Model Context Protocol standardized how an assistant reaches external tools and data [23]. A rapidly growing literature has explored what happens when this capability is pointed at biological data analysis: autonomous multi-omics agents [24], agentic frameworks for biomedical data science [25], surveys of agent architectures across genomics, proteomics and spatial biology [26,27], and the first serious benchmarks of end-to-end analysis competence [28].

Most of that work asks the assistant to write the analysis: to generate, for example, a pandas or Seurat script from scratch, execute it, and debug it. This is a powerful approach, but for a mature and well-specified method it is arguably not the most useful division of labor. Code generated on the fly is not reviewed, not versioned, not tested, and not the same twice; its statistical choices are whatever the model happened to sample. For a well-specified analysis with known failure modes, the last thing one wants is a fresh implementation each time.

The alternative is to keep the implementation fixed and audited, and to let the assistant supply what it is actually good at: understanding an unstructured request, inspecting messy inputs, asking the clarifying question, selecting among a bounded set of validated operations, and explaining the result. This is precisely the role of an agent skill — a self-contained package consisting of a structured instruction file (`SKILL.md`), optional executable scripts, and reference documents, which the assistant discovers by name and description, loads when relevant, and follows [29]. The architecture is progressively disclosed: a few dozen tokens of metadata sit in context permanently; the full procedure loads only when the task matches; deep reference material loads only when a specific sub-question demands it [29]. Xu and Yan describe the design as “an onboarding guide for a new hire” — comprehensive institutional knowledge that stays out of the way until it is needed. This suggests a modest reframing, and it is the one around this paper is organized: the unit of scientific software distribution may usefully be extended from the program to the program together with the knowledge of how to use it. A package is a set of capabilities. A skill is a set of capabilities plus the judgement about when to invoke them, what to check first, and what the answer is allowed to mean — the tacit knowledge that typically lives in the head of the group’s bioinformatician and leaves with them. Writing that knowledge down in a form a machine can follow has, we think, some of the same value as publishing a protocol.

We therefore present MErlin, whose nature is dual. As a package, it is a seventeen-module command-line toolkit for prokaryotic methylation multi-omics that works perfectly well from a terminal. As a skill it is a document that teaches an assistant to drive that toolkit the way a careful bioinformatician would, including the parts where the careful bioinformatician says no.

We describe the architecture (please refer to the paragraphs from 2.1 to 2.5), the skill layer and what it actually enforces (§2.6), validation on synthetic data with planted ground truth (§2.7), and a brief real-world illustration in the Antarctic marine bacterium *Pseudoalteromonas haloplanktis* TAC125 [30] in which the conversational interface returns a correctly qualified result (§2.8). In Section 3 we then discuss the obligations this mode of distribution creates — chiefly around reproducibility, biological interpretation and overclaiming.

## 2. Results

### 2.1 A two-body tool

MErlin is distributed as two artefacts with a deliberate division of labour.

The package is a Python ≥`3.10` library exposing a single console entry point, `merlin`, with seventeen subcommands. It has a small hard dependency set (`numpy`, `pandas`, `scipy`, `matplotlib`) and a set of optional extras, each of which degrades to a stated fallback rather than failing: without `cooler`, a native HDF5 reader handles .cool files; without `modkit`, an internal `pysam pileup` runs; without `R`, the `circos` figure is drawn in `matplotlib`. `merlin doctor` reports, for every module, whether it is ready, degraded or unavailable, and for the latter two, why, naming the package to install. This matters more than it sounds: installability is a documented and severe problem in the omics tool ecosystem [31], and a tool that cannot report its own state cannot be driven by an agent.

The skill is a directory (`skill/merlin/`) containing `SKILL.md`, a `SETUP.md`, a `references/tree` (`tools`, `statistics`, `preflight`, `interpretation`, `troubleshooting`) and a `scripts/directory`. It contains no analysis logic. It is procedural knowledge about the package, structured for progressive disclosure [29], and it is versioned and reviewed in the same repository as the code it describes.

Neither is sufficient alone in the intended use case. The skill without the package lets an assistant explain MErlin; the package without the skill is a competent CLI that presumes a competent CLI user.

### 2.2 Seventeen modules over six data layers

The modules divide into analysis, integration and orchestration (**Figure 1, Table 1**).

**Table 1.**
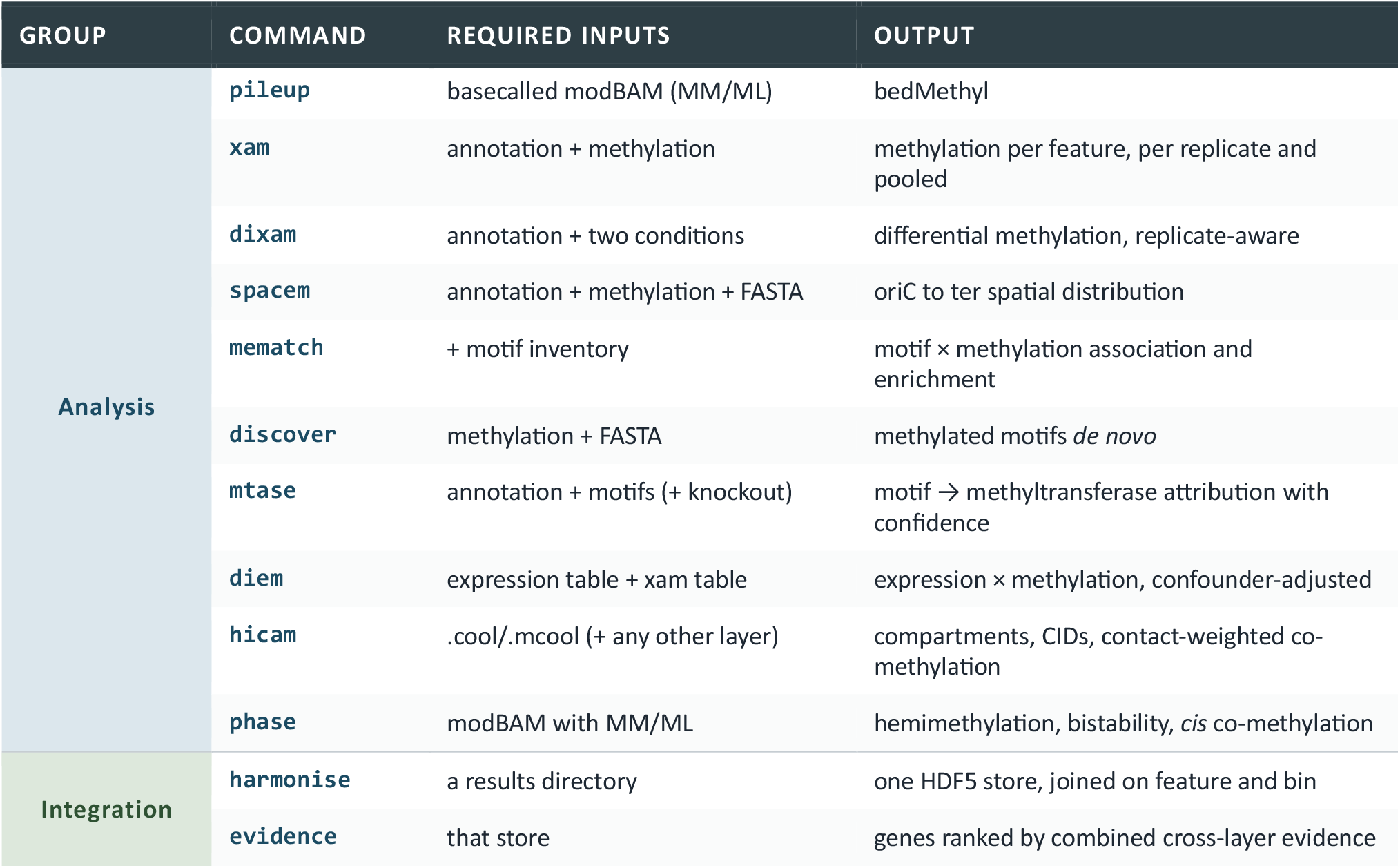
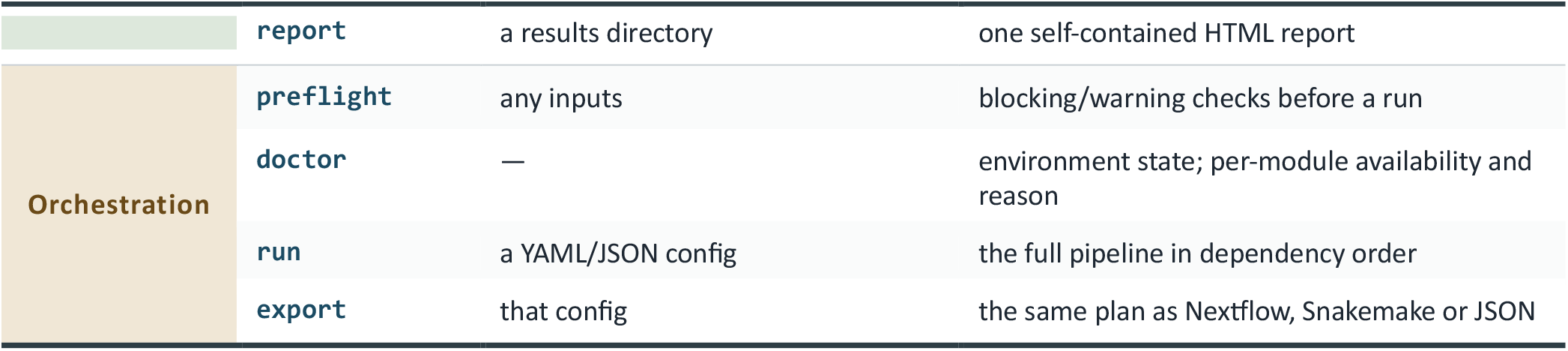
The seventeen MErlin modules, by role. Every module runs on whatever inputs exist; a module whose required inputs are absent is skipped and named in the merlin run --dry-run execution plan.

| GROUP | COMMAND | REQUIRED INPUTS | OUTPUT |
| --- | --- | --- | --- |
| Analysis | pileup | basecalled modBAM (MM/ML) | bedMethyl |
|  | xam | annotation + methylation | methylation per feature, per replicate and pooled |
|  | dixam | annotation + two conditions | differential methylation, replicate-aware |
|  | spacem | annotation + methylation + FASTA | oriC to ter spatial distribution |
|  | mematch | + motif inventory | motif × methylation association and enrichment |
|  | discover | methylation + FASTA | methylated motifs *de novo* |
|  | mtase | annotation + motifs (+ knockout) | motif → methyltransferase attribution with confidence |
|  | diem | expression table + xam table | expression × methylation, confounder-adjusted |
|  | hicam | .cool/.mcool (+ any other layer) | compartments, CIDs, contact-weighted co-methylation |
|  | phase | modBAM with MM/ML | hemimethylation, bistability, *cis* co-methylation |
| Integration | harmonise | a results directory | one HDF5 store, joined on feature and bin |
|  | evidence | that store | genes ranked by combined cross-layer evidence |
|  | **report** | a results directory | one self-contained HTML report |
| **Orchestration** | **preflight** | any inputs | blocking/warning checks before a run |
|  | **doctor** | — | environment state; per-module availability and reason |
|  | **run** | a YAML/JSON config | the full pipeline in dependency order |
|  | **export** | that config | the same plan as Nextflow, Snakemake or JSON |

**Figure 1.**
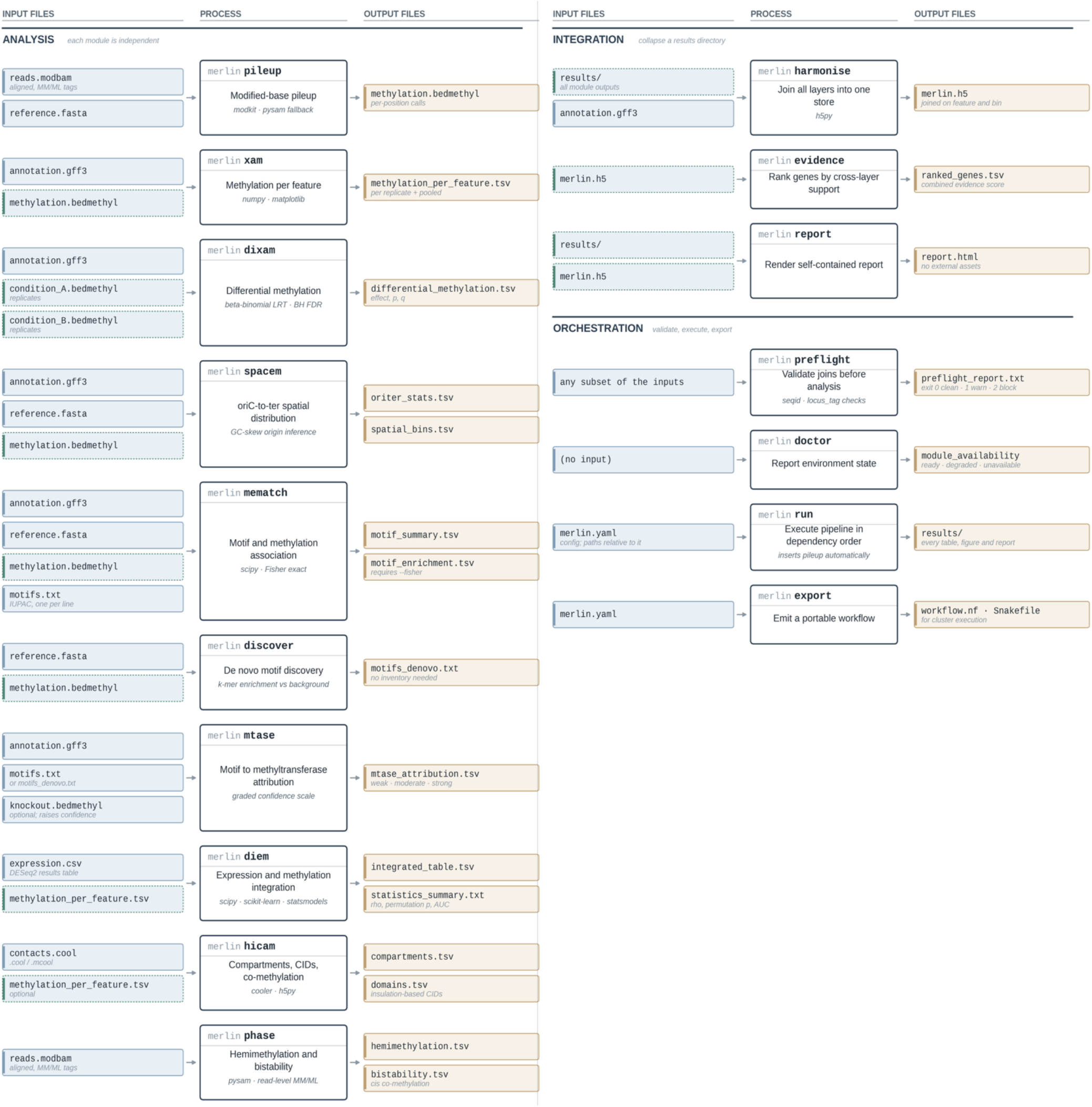
MErlin workflow. Seventeen modules over six data layers. (a) Ten analysis modules, each running on whatever inputs exist. (b) Integration into one joined store, gene ranking and HTML report; orchestration for validation, execution and workflow export. Blue: user-supplied input; dashed blue: produced by an upstream module; amber: module output. Italic text names the method or dependency used. Every module writes a provenance sidecar. `preflight` runs first because both identifier joins fail silently rather than raising.

#### Analysis

`pileup` converts basecalled modBAM (MM/ML tags) to bedMethyl, using `modkit` where available and an internal `pysam` backend otherwise.

`xam` computes methylation per annotated feature, per replicate and pooled, with configurable sub-feature regions (upstream windows, gene body).

`dixam` performs replicate-aware differential methylation between two conditions.

`spacem` places methylation on the oriC-ter axis, with GC-skew-based origin inference and an explicit override.

`mematch` tests association and enrichment between a supplied motif inventory and observed methylation.

`discover` finds methylated motifs de novo when no inventory exists.

`mtase` attributes a motif to a methyltransferase gene, with a confidence level that rises from annotation only to knockout-supported. `diem` relates expression to methylation with confounder adjustment.

`hicam` calls compartments and chromosomal interaction domains from `.cool`/.`mcool` matrices and computes contact-weighted co-methylation. phase works at read level to quantify hemimethylation, bistability and cis co-methylation.

#### Integration

`harmonise` collapses a results directory into a single HDF5 store with every table joined on feature and on genomic bin. evidence ranks genes by combined support across whichever layers are present. `report` emits one self-contained HTML file.

#### Orchestration

`preflight` runs blocking and warning checks on any set of inputs (§2.3).

`doctor` reports environment state. run executes a YAML/JSON-configured pipeline in correct dependency order, inserting `pileup` steps automatically when the config supplies modBAMs rather than bedMethyl.

`export` emits the same plan as a Nextflow [32] or Snakemake [33] workflow for cluster execution.

The governing rule is that every module runs on whatever inputs exist. A methylation-only project — annotation plus bedMethyl, nothing else — is a first-class case that yields real answers from `xam`, `dixam`, `discover`, `mematch`, `spacem` and `phase`. `merlin run --dry-run` prints the execution plan with every skipped module and the specific missing input that caused the skip. A plan with three skips is a normal methylation-only run, not a broken configuration, and the tool says so.

### 2.3 Preflight: exposing silent integration failures

`merlin preflight` accepts any subset of the inputs and returns `exit code 0` (clean), `1` (warnings) or `2` (blocking). It checks, among other things:

– seqid namespace agreement across FASTA, GFF3, bedMethyl and Hi-C matrix. Version-suffix mismatches (NZ_XXX.1 vs NZ_XXX) are reconciled automatically and reported; genuine namespace disjunctions (e.g., accession vs. contig name) are blocking.
– locus_tag agreement between the annotation and the expression table.
– Coordinate-system sanity: methylation positions falling outside the declared contig lengths.
– Coverage distribution in the methylation calls, which is where the difference between a depth column and a methylated-read count becomes visible (§2.8).
– Hi-C bin size versus gene size, which determines whether a hicam result can speak about genes at all.

This module exists because of the eventual failure mode described in §1.2, and it is one of the main practical reasons to run MErlin rather than write the joins by hand. The skill treats preflight as mandatory: an assistant following it will not proceed to analysis on a blocking exit code and will surface the specific mismatch to the user in plain language.

### 2.4 Experimental design bounds the licensed inference

MErlin’s statistical modules are constrained by the experimental design they receive, and they say so in their output.

`dixam` with biological replicates per condition fits a replicate-aware beta-binomial model and reports a likelihood-ratio test with Benjamini–Hochberg control of the false discovery rate [34]. `dixam` with one sample per condition runs a within-sample test on read counts and labels the result n_replicates = 1v1. That p-value is a statement about the sampling of reads at a locus; it is not a statement about biological reproducibility, and it does not by itself constitute evidence for differential methylation between conditions. MErlin records the distinction in the output table, in the report, and in the provenance sidecar, and the skill is instructed to reproduce it verbatim in any written summary. Providing replicates is the only thing that changes it.

Two further design traps are handled explicitly because they invert conclusions while looking entirely normal:

– `--labels` is positional. In `dixam` it defaults to control treatment irrespective of file contents. A swapped pair inverts every differential call and produces output indistinguishable from a correct run. The same applies to the sign convention of an imported DESeq2 [20] log2FoldChange column: nothing in the file records which condition was the numerator. The skill’s first-order rule is to ask the user which condition is the treatment, in both layers, before running, and to refuse to guess.
– `xam --include-zero`. Without it, features with no methylation call are absent from the table rather than present with value zero, so any downstream correlation in diem silently describes only the methylated subset. The skill enforces `--include-zero` whenever `diem` is in the plan.

`mtase` implements a graded confidence scale rather than a binary call: attribution from annotation alone is weak; supported by motif–gene co-occurrence, moderate; supported by loss of the motif in a methyltransferase knockout, strong. Only the last licenses the sentence “motif X is methylated by gene Y”, and it is the only condition under which MErlin’s interpretive guidance permits causal language anywhere in the toolkit.

### 2.5 Provenance

Every module writes a `<prefix>.provenance.txt` sidecar next to its tables recording the full command line, the version of MErlin and of every relevant dependency, and the SHA-256 of each input file. This is what a methods section cites. In the conversational mode it also serves a second purpose: it is the audit trail that converts a natural-language session into something a reviewer can check, because the provenance file records what was actually executed regardless of how the request was phrased (§3.3).

### 2.6 The skill layer

The skill is what makes the following interaction possible, and it is worth being precise about what it does and does not do. A user with no command-line experience uploads a folder of bedMethyl files and a Prokka GFF3 to a Claude session with the MErlin skill enabled, and writes:

> *I have bedMethyl for a Sinorhizobium wild type and a dam knockout, three replicates each, plus the Prokka annotation. What can MErlin tell me?*

A correctly loaded skill produces a specific sequence: it runs `merlin doctor`, then `merlin preflight` on the supplied files; it reports the namespace state; it identifies the design as replicated two-condition and therefore eligible for the beta-binomial `dixam`; it notes that a knockout condition upgrades mtase attribution from weak to strong; and before running anything, it asks which condition is the treatment. If it instead offers to write a `pandas` script, the skill did not load, and this is the documented smoke test for a correct installation.

What the skill contributes is therefore not autonomy but constraint. Its content is, in rough proportion: a decision procedure mapping available inputs to eligible modules; a set of mandatory pre-run checks; a list of questions that must be answered by the human before execution; the design to licensed-claim mapping of §2.4; and an interpretation guide whose main function is to prohibit sentences. The `references/interpretation.md` file, for instance, exists chiefly to enforce the distinction between “methylation co-varies with expression” (a finding MErlin can support) and “methylation regulates expression” (which it cannot, at any p-value), and to require that a 1v1 design be described as screening candidates rather than establishing differences. None of this knowledge is new. All of it is the ordinary practice of a careful analyst. What is added here is that it is written down, shipped with the code, version-controlled alongside it, and applied identically on every run by a user who has never met the analyst.

### 2.7 Validation against synthetic ground truth

MErlin ships a synthetic dataset (`testdata/`, regenerable with `tools/make_testdata.py`) constructed with known answers planted in it, and a `pytest` suite that asserts the biology comes back out. The planted truth comprises:

– three methyltransferase systems with distinct recognition motifs — GATC (6mA), GANTC (6mA) and CCWGG (5mC) — at specified methylation frequencies;
– a set of hypomethylated-and-downregulated genes, i.e., a true negative methylation– expression association of known effect size;
– Hi-C contact domains with defined boundaries;
– a bistable locus with a defined mixture of methylated and unmethylated reads;
– a deliberate seqid version-suffix mismatch between FASTA and GFF3.

The suit asserts that:

– `discover` recovers all three motifs de novo without being given a motif list;
– `mematch` assigns them the expected enrichment;
– `mtase` attributes each to the correct planted gene;
– `dixam` recovers the planted differential features;
– `diem` recovers the planted correlation with the correct sign;
– `hicam` recovers the domain boundaries;
– `phase` identifies the bistable locus and no others;
– `preflight` detects and reconciles the seeded mismatch.

A parallel configuration (`merlin.from_bam.yaml`) executes the identical run starting from a synthetic modBAM with no bedMethyl anywhere, verifying the `pileup` path.

### 2.8 Real-data illustration: methylome and transcriptome of the bacterium P. haloplanktis TAC125 grown in two media

To show the conversational mode on real data, we summarise a comparison of the methylome and transcriptome of *P. haloplanktis* TAC125 [30] between two growth media (here MA and TYP). The entire analysis was specified in natural language; no merlin command was typed by the user.

Starting material was unaligned PacBio HiFi modBAM at ∼1,840× and ∼2,130× coverage, systematically subsampled to 263× and 266× (MM/ML tags retained), aligned and passed to `merlin pileup`. Both conditions carry the same methylome. `discover` identified a single Dam-type GATC 6mA system de novo (∼15,500 sites; 91.6% of assayed GATC calls methylated), symmetric on both strands; `mtase` attributed it to the annotated dam gene at confidence = weak (annotation only), correctly declining to make the attribution stronger in the absence of a knockout. `phase` found zero hemimethylated and zero bistable loci in either condition. `dixam` returned no differentially methylated 6mA feature (minimum q = 1.0). 5mC was sparse (mean 4.3–4.5% modified), depleted on CG relative to background (site-level OR = 0.35), and produced only five candidate differences.

`diem` found a weak negative relationship between methylation density and the TYP-vs-MA expression log_2_ fold change (Spearman ρ = ™0.087, permutation p = 2 × 10^−4^, n = 2,419), essentially unchanged after confounder adjustment (partial ρ = ™0.083); a random-forest classifier reached a cross-validated AUC of 0.55 against a differential-expression prevalence of 0.135 — i.e., negligible predictive value. Differential methylation against differential expression was not significant (ρ = +0.013, p = 0.56).

The conclusion MErlin reported, is that the MA vs TYP transcriptional response in this dataset is not accompanied by a change in DNA methylation. These data were already tested running MErlin as toolkit, and the agent skill confirmed our previous results.

Biologically this is the expected shape of the result rather than a surprise, and it is worth setting out why, since the reasoning is what the layered analysis buys. Dam-type systems in *gammaproteobacteria* methylate GATC to near-saturation and act constitutively; the documented cases in which GATC methylation controls transcription are local and depend on a DNA-binding protein occluding a particular site so that it escapes methylation for at least part of the cell cycle [3,6]. Such a mechanism leaves a signature: a site that is persistently hemimethylated or bimodally methylated across the population. phase found none, in either condition, and dixam found no condition-dependent change in the 6mA layer. The two routes by which this system could plausibly carry a condition-responsive regulatory signal in these data are therefore both closed, which is a stronger statement than the absence of a correlation on its own would support. The 5mC layer is not interpreted further here: at a mean modified fraction of 4.3–4.5% it sits close to the level at which single-molecule 5mC calls in bacteria are of uncertain reliability, and the five candidate differences should be treated as unresolved rather than as a weak positive. The residual weak negative association recovered by diem is of the magnitude expected from gene-level composition — GATC density co-varies with gene length and GC content — and is essentially unchanged by adjustment for those covariates, which is consistent with a compositional rather than a regulatory origin. One design caveat should be recorded alongside the three the tool raised. MA and TYP differ in growth rate as well as in composition, and the fraction of cells carrying an active replication fork sets the genome-wide level of hemimethylation; the two media are therefore not matched on the variable most likely to move the measurement. That the observed hemimethylation was zero in both conditions makes this unlikely to have driven the result here, but a comparison of media should in general be harvested at matched growth phase. What these data cannot exclude is a small number of individually regulated promoters whose effect is invisible to a genome-wide statistic; resolving that would require a dam mutant and site-level analysis of the promoters concerned, which is precisely the experiment that mtase’s weak confidence label points to.

Three caveats were raised by the tool without prompting, and they illustrate why we include this example. First, the design is one sample per condition; `dixam` therefore ran its single-sample test and its q-values screen candidates rather than establishing differential methylation. Second, PacBio’s A+a. MM list is sparse (only candidate adenines are enumerated) so the 6mA bedMethyl “coverage” column counts methylated reads rather than depth, and `mean_percent_modified` is 100 by construction; `preflight`’s coverage-distribution check surfaced this, and the 6mA layer was consequently treated as a count/presence track (dixam --test-on calls, --min-coverage 20, `--min-frac 0.5`). Third, GC skew was too flat on all three main replicons (relative amplitude 0.010–0.012) for reliable origin inference, so every `spacem` ori to ter statistic was flagged unreliable rather than reported.

The negative appears to be the correct answer here, and arriving at it required the kind of skepticism that the skill encodes.

## 3. Discussion

### 3.2 What changes when the interface is conversational

MErlin’s contribution does not rest on novelty in the underlying algorithms as several modules intentionally implement established statistical approaches. Rather, it resides in whether coupling fixed, inspectable software to explicit procedural guidance can make complex multi-omics workflows easier to operate without relaxing the conditions required for valid inference. The *P. haloplanktis* TAC125 analysis of §2.8 illustrates what the conventional mode demands: the user must know that PacBio’s sparse MM encoding gives the coverage column a different meaning from a nanopore bedMethyl; that dixam must therefore test on calls rather than fractions; that a flat GC skew invalidates origin inference; that a 1v1 design licenses no differential claim; and that a permutation p of 2 × 10^−4^ on a ρ of ™0.087 is a statement about the null distribution, not about biological importance. Every one of these is knowable. None of them is in a `--help` string. Collectively they can be the difference between a correctly reported negative and an overstated positive.

The conventional answer is that this is why laboratories employ bioinformaticians, and it is a good answer where a bioinformatician is employed. It fails at the long tail: the microbiology group with one nanopore run and no computational staff, the student between rotations, the collaborator who needs the methylation layer of someone else’s project analyzed in short time. That long tail is where the 89% figure of Barone et al. [21] lives, and it is a consistent fraction of world-wide science.

An assistant equipped with a skill does not replace a bioinformatician. What it does is make the checklist portion of expertise — the part that is procedural, enumerable, and most often skipped under time pressure — available to everyone, on every run, without fatigue. We take this to be a limited claim, and a defensible one.

### 3.2 Why constraint matters in conversational analysis

The obvious objection to conversational analysis is that language models are agreeable. Conversational systems can follow a user’s framing too readily, particularly when the requested biological narrative is stronger than the experimental design supports. When this behavior is combined with the flexibility of on-the-fly code generation, this is a combination that favors false positives.

MErlin’s response is architectural. Because the assistant selects among a fixed set of audited modules rather than writing analysis code, it cannot quietly choose a more permissive test; because the statistical claim is bound to the design at the level of the module’s own output (n_replicates = 1v1 is printed by MErlin, not asserted by the assistant), the constraint survives an enthusiastic user; and because the interpretation guidance is written as a set of prohibited sentences rather than encouraged ones, the failure mode is under-rather than over-claiming. In the *P. haloplanktis* TAC125 run, the tool volunteered three reasons to distrust its own output. We therefore propose a general design principle for scientific agent skills: a useful skill should constrain the action and inference space of the assistant at least as much as it expands access to the software. A skill that makes an assistant more capable of arbitrary analysis is a liability. A skill that makes it reliably refuse to run before checking a join is an asset. The current benchmarking literature for bioinformatics agents [28] measures task completion; we would argue that appropriate refusal deserves equal weight, and that a benchmark on which the highest score requires declining several of the tasks would be a better instrument.

### 3.3 Reproducibility of a conversation

A natural-language session is not itself reproducible: the same request phrased twice may produce different intermediate reasoning, and the model behind the assistant is updated on a schedule the user does not control. This is a genuine problem and we do not want to minimize it.

MErlin’s position is that the conversation is the interface, not the record. What is reproducible is what was executed, and that is captured in three places: the provenance.txt sidecars (command line, versions, input SHA-256), the YAML config that merlin run consumes, and the workflow exported by merlin export for Nextflow [32] or Snakemake [33]. A session that produced a figure can therefore be reduced to a config file and re-run deterministically by anyone, including on a cluster, with no assistant involved.

In this model, natural language is used to arrive at an appropriate analysis, whereas the configuration and provenance files record what was done. For publication, we therefore recommend archiving the executable configuration, software release and provenance records together with the data required to reproduce the reported results; the conversational transcript may be informative, but it is not a substitute for these artefacts.

### 3.4 Limitations

MErlin has several limitations that should define the scope of the present claims. First, the agent skill cannot guarantee correct behavior: it remains dependent on the capabilities and instruction-following behavior of the host model and agent environment. The skill should therefore be evaluated empirically and versioned together with the software configuration used in a study. Second, constraining the assistant to audited modules transfers responsibility to the modules themselves; implementation errors, inappropriate default parameters or unrecognized biological assumptions in the package remain possible and require conventional software testing and expert review. Third, the current real-data illustration contains one methylation sample per condition and therefore cannot establish biological reproducibility of differential methylation. Finally, portability of the skill across multiple LLM providers or agent frameworks should not be assumed without direct testing. These limitations motivate the synthetic software benchmark reported here, and the dedicated evaluation of the agent layer set out in §3.5.

### 3.5 Evaluating the agent layer

The claim that the skill changes what a non-expert obtains is, at present, supported by one worked example. It is a testable claim, and the synthetic dataset of §2.7 already provides the instrument, since the traps it contains are objectively scorable. We therefore set out the evaluation we consider necessary, which we intend to implement.

The comparison requires three arms receiving identical inputs and prompts:

1. an assistant with code execution but without MErlin, which must write its own analysis;
2. an assistant with the MErlin package installed but without the skill, which must discover the interface from `--help` and the README;
3. an assistant with both.

The middle arm is the informative one, since without it the contribution of the package cannot be separated from the contribution of the procedural knowledge, and it is the latter that this paper is about. Endpoints should not be restricted to task completion, for the reason given in §3.2.

Three are available:

– recovery of the planted ground truth;
– detection of the seeded traps, each of which is pass/fail — the seqid version mismatch, the swapped `--labels` and DESeq2 sign conventions, a 1v1 design reported as differential, an omitted `--include-zero`, a sparse-MM coverage column, and an unreliable GC-skew origin call;
– the calibration of the written summary, which requires a rubric fixed in advance and blinded grading.

Whether a re-runnable configuration and provenance record is produced at all is a fourth, and the arm without MErlin will usually fail it by construction.

The same request, phrased the same way, does not produce the same session twice: the assistant may ask a different clarifying question, or reach the same answer by a different route since sessions are not deterministic; each arm has to be repeated and its variability reported rather than a single run.

### 3.6 Outlook

A large number of bacterial methylomes have been deposited and described but only very few have been interrogated against transcription, motif complement or population structure, and the comparative question — which methyltransferases have phenotypes, in which lineages, and under which conditions — remains largely open [14]. Lowering the interface cost can broaden research on the biology of DNA methylation in prokaryotes. Extension beyond prokaryotes is possible, but it is worth being explicit about which parts transfer.

Format-level components move unchanged: modBAM MM/ML tags are identical for eukaryotic long-read data, so `pileup` applies directly, and `preflight`, `harmonise`, `report`, `run` and `export` are domain-agnostic. Two analytical modules arguably fit eukaryotic data better than bacterial: `hicam`, since compartments and interaction domains are eukaryotic concepts imported into bacteria, and `phase`, whose read-level machinery maps onto allele-specific methylation and imprinting. A second group requires the unit of analysis to be redefined before it means anything: `xam`, `dixam` and `diem` aggregate per gene, whereas eukaryotic methylation must be summarised over CpG islands, promoters, enhancers, gene bodies and repeat classes — a distinction that is biological rather than cosmetic, since promoter and gene-body methylation associate with expression in opposite directions, and differential calling requires smoothing across neighbouring CpGs rather than independent per-feature tests. A third group is prokaryote-specific by construction: `discover`, `mematch` and `mtase` assume that methylation is motif-driven and attributable to a sequence-specific methyltransferase, which has no counterpart in chromatin-directed DNMT targeting, and `spacem` would have to be replaced by a replication-timing module rather than adapted, even though the confounder it addresses persists in the form of late-replicating partially methylated domains. Two further obstacles are practical: genome size makes the current in-memory design inadequate above the prokaryotic scale, and cell-type composition introduces a dominant confounder with no bacterial analogue. We would therefore claim portability for the skill layer more confidently than for the package: the mapping from experimental design to licensed inference is domain-general, and eukaryotic epigenomics has its own catalogue of silent failures — genome-build and identifier mismatches, the promoter/gene-body sign inversion, composition effects — of exactly the kind this approach is meant to make loud.

## 4. Methods

### 4.1 Implementation and availability

MErlin is written in Python (≥`3.10`). Required dependencies are `numpy`, `pandas`, `scipy` and `matplotlib`. Optional extras are `ml` (`scikit-learn`, `statsmodels`, `seaborn`), `store`/`hic` (`h5py`), `cooler` [35], `reads` (`pysam`) and `config` (`pyyaml`); each absent extra disables specific functionality with a stated reason reported by merlin doctor rather than raising at run time. Optional external binaries are `modkit` (Oxford Nanopore Technologies), `minimap2`/`samtools` for aligning unaligned modBAM, and `R` with `optparse`, `data.table`, `circlize` and `Biostrings` for the `roundtable.R circos` figure; each has an internal fallback. A `conda` environment file is provided.

Installation is `conda env create -f environment.yml` followed by `pip install -e “.[all]”`. MErlin is not currently on `PyPI`; release wheels are published on GitHub.

### 4.2 Module methods

`pileup`. modBAM MM/ML tags are converted to per-position modified-base counts. The default backend is modkit where present; otherwise an internal `pysam` implementation applies the same modification-probability threshold `(--filter-threshold`, `default 0.7`), minimum mapping quality `(--min-mapq`) and strand handling, and emits bedMethyl.

`xam`. Per-feature methylation is computed by intersecting bedMethyl positions with annotation features on seqid, optionally restricted to sub-feature regions specified as upstream: -300, body, etc. Values are reported per replicate and pooled. `--include-zero` retains features with no methylation call as explicit zeros.

`dixam`. With ≥2 replicates per condition, methylated and unmethylated read counts per feature are modelled as beta-binomial and compared by likelihood-ratio test, with Benjamini–Hochberg FDR control [34]. With one sample per condition, a within-sample count test is applied and the design is recorded as n_replicates = 1v1 in the output and provenance. `--test-on` calls switches the response from methylated fraction to methylated-call count for sparse-MM inputs (§2.8). `--labels` assigns condition names positionally, defaulting to control treatment.

`spacem`. Origin and terminus are inferred from cumulative GC skew unless supplied explicitly with `- -oriC/--ter`; the relative skew amplitude is reported and results are flagged unreliable below a threshold. Methylation is binned along the normalised ori to ter axis for each replichore.

`mematch`. Observed methylation is tested for association with a supplied motif inventory at site level (odds ratios against a background of unmethylated matching sites) and at feature level (enrichment of motif-containing features among methylated features).

`discover`. Methylated positions are extracted with flanking sequence from the reference and enriched k-mers/degenerate motifs are identified against a background of unmethylated positions of the same base, with no motif list supplied.

`mtase`. Candidate methyltransferase genes are identified from the annotation and matched to discovered or supplied motifs; a confidence level is assigned (weak — annotation only; moderate — supported by co-occurrence; strong — supported by loss of the motif in a supplied knockout condition).

`diem`. An expression table (e.g., DESeq2 results [20]) is joined to the `xam` table on locus_tag. Association between methylation density and expression (or log_2_ fold change) is quantified by Spearman correlation with a permutation null, and by partial correlation adjusting for declared confounders (gene length, GC content, replicon, expression level). A random-forest classifier of differential-expression status from methylation features is evaluated by cross-validated AUC against the observed DE prevalence.

`hicam`. `.cool`/`.mcool` matrices are read via `cooler` [35] or a native HDF5 reader. Compartments are called from the first eigenvector of the observed/expected correlation matrix; chromosomal interaction domains [17,18] are called from an insulation profile. Contact-weighted co-methylation is computed as the contact-frequency-weighted correlation of bin-level methylation.

`phase`. Read-level MM/ML tags are used to score, per site, the fraction of reads methylated on each strand (hemimethylation), the bimodality of the per-read methylation distribution (bistability), and pairwise cis co-methylation between sites spanned by the same read.

`harmonise`, `evidence`, `report`. `harmonise` writes all module outputs into one HDF5 store joined on feature and on genomic bin. `evidence` ranks genes by combined support across available layers. `report` renders a self-contained HTML document with no external assets.

### 4.3 Synthetic benchmark

The synthetic dataset is generated by `tools/make_testdata.py` with a fixed random seed. It comprises a synthetic circular replicon with an annotation, three methyltransferase systems (GATC 6mA, GANTC 6mA, CCWGG 5mC) applied at defined frequencies, an expression table containing a planted set of hypomethylated-and-downregulated genes, a Hi-C matrix with defined domain boundaries, a bistable locus with a defined methylated-read mixture, a synthetic modBAM carrying MM/ML tags consistent with the bedMethyl, and a deliberate seqid version-suffix mismatch between FASTA and GFF3. The `pytest` suite in `tests/` asserts recovery of each planted feature. Reproduce with: `python3 tools/make_testdata.py`

### 4.4 *P. haloplanktis* TAC125 data

Basecalled PacBio HiFi modBAM for P. haloplanktis TAC125 [30] in two growth conditions (MA, TYP) carried `A+a.`, `T-a.` and `C+m?` MM entries at approximately 1,840× and 2,130× coverage. Reads were systematically subsampled to 263× and 266×, discarding kinetics tags and base qualities while retaining MM/ML, then converted and aligned with `samtools fastq -T MM,ML | minimap2 -ax map-hifi -y | samtools sort`. `merlin pileup` was run with the internal `pysam` backend (`modkit` could not be installed in the execution environment). Because PacBio’s `A+a.` list enumerates only candidate adenines, the 6mA layer was treated as a count/presence track: `dixam --test-on calls --min-coverage 20 --min-frac 0.5`. Differential expression was supplied as a DESeq2 [20] results table with positive log_2_ fold change meaning up-regulated in TYP (confirmed with the data provider); `dixam --labels MA TYP` was used so that the methylation effect is likewise TYP ™ MA. Analysis was driven end to end through natural-language requests to a Claude assistant with the MErlin skill enabled; provenance sidecars for every step are deposited with the results.

### 4.5 Skill implementation

The skill follows the Agent Skills convention [29]: a directory whose name matches the name field of the YAML frontmatter in its `SKILL.md`, containing that file plus `SETUP.md`, `references/` and `scripts/`. Progressive disclosure is used throughout: `SKILL.md` contains the decision procedure and mandatory checks; the detailed statistical and interpretive material lives in `references/statistics.md`, `references/interpretation.md`, `references/preflight.md`, `references/tools.md` and `references/troubleshooting.md`, which are loaded only when the relevant question arises. Installation is either by uploading a `.s`kill archive of the folder through the Claude interface, or by copying the folder to `∼/.claude/skills/` (`personal)` or `.claude/skills/ (project)`. Correct installation is verified with the smoke test described in §2.6.

## Data and code availability

MErlin is open source and available at https://github.com/IacopoPasseri/MErlin, including the package, the agent skill, worked example configurations, the synthetic test dataset and its generator, and the test suite. The version described here is 2.1.0.

## Competing interests

The authors declare no competing interests.

Note: no author has a financial relationship with Anthropic or any other provider of the assistant software discussed.

## Notes

### Competing Interest Statement

The authors have declared no competing interest.

